# A simple, cost efficient assay for assessing the functional impact of single and multi-gene variant combinations

**DOI:** 10.1101/2025.07.11.663622

**Authors:** Mats Georg Telge, Ioannis Evangelakos, Christian Schlein

## Abstract

Obesity is a major global health burden driven by both genetic and environmental factors. While monogenic forms are rare, their identification is essential given the availability of targeted therapies. However, genetic testing frequenctly uncovers many variants of unknown significance (VUS), highlighting the need for functional assays to enable precision therapeutic decisions. Here, we describe a simple and cost-effective *in vitro* assay enabling high-throughput functional analysis of genetic variants, including combinations across different genes. As a proof of concept, we applied this platform to systematically assess the most common *LEP* and *LEPR* variants based on gnomAD v2.1.1 allele frequencies and benchmarked them against known pathogenic controls. In total, we assessed 35 *LEP* variants (including 3 controls) and 30 *LEPR* variants (including 3 controls), measuring in total 2,100 unique variant combinations. This approach identified 19 VUS to be putatively pathogenic. Importantly, 806 combinations of *LEP* and *LEPR* variants coexpressed with their respective wild type allele exceeded pathogenicity thresholds, revealing a potential digenic mutational burden. In sum, this assay offers a scalable strategy for the functional characterization of obesity-associated variants and offers a valuable tool for on-demand VUS interpretation in clinical diagnostics.

## Introduction

Obesity represents one of the most significant public health challenges of the century, with profound implications for global morbidity, mortality, and economics in healthcare systems^1,2^. While lifestyle and environmental factors are major contributors^3^, a substantial body of evidence highlights the critical role of genetic predisposition in the development and maintenance of obesity^4,5^. Among the predominantly genetically influenced forms, rare monogenic variants often affect key hypothalamic pathways involved in appetite regulation and energy homeostasis and have been identified in a subset of individuals mostly with severe, early-onset obesity^6^. Although these monogenic forms account for only a small fraction of obesity cases^7^, their accurate diagnosis is of high clinical relevance, as specific targeted therapies, such as melanocortin-4 receptor (MC4R) agonists, have recently become available^8,9^. However, routine genetic testing frequently uncovers variants of uncertain significance (VUS)^10,11^. Functional characterization of these variants is therefore essential to enable molecular diagnosis and guide individualized treatment strategies.

One of the best-studied pathways in the regulation of food intake is the Leptin-Leptin receptor pathway^12^. Leptin, encoded by *LEP,* is secreted by adipocytes and acts via the Leptin receptor, encoded by *LEPR*, in the nucleus arcuatus^13,14^. In obese individuals, leptin resistance - leading to increased food intake, insulin resistance, and dyslipidemia^4^ - is common and often influenced by variants in *LEP* and *LEPR*^5^ or in genes functionally related to them. This highlights the importance of a detailed understanding of the mechanisms underlying energy homeostasis in humans.

Over 200 cases of LEP or LEPR deficiency have been reported^6,7^ since the first description of diseases associated to biallelic pathogenic variants in these genes, presenting with severe early-onset obesity, hyperphagia, immune dysfunction, delayed puberty, reduced linear growth, pituitary dysfunction, and hypogonadotropic hypogonadism^15,16^.

Heterozygous *LEP* and *LEPR* variants are known to modestly increase BMI and body fat percentage^17^. However, how compound heterozygosity in each gene and digenic interactions between them contribute to obesity remains unclear.

The master regulator of central appetite control downstreaam of leptin signalling pathway is signal transducer and activator of transcription 3 (STAT3), mainly by driving transcription of *POMC*^18^. For this reason, the current gold standard approach for assessing the functionality of *LEP* and *LEPR* variants involves measuring their effects on STAT3 phosphorylation *in vitro* using Western blot^19^ or analyzing patient data^20^. However, this method is insufficient for high-throughput evaluation, particularly given the exponential rise in possible variant combinations and that LEPR dimerization upon leptin binding activates multiple downstream pathways^21^.

To address this, we aimed to comprehensively characterize the most common *LEP* and *LEPR* variants by developing an assay capable of measuring heterozygous-like, compound-heterozygous-like, and digenic variant combinations, which can be performed cost-efficiently using standard laboratory equipment. For this purpose, we developed and optimized a dot blot protocol for phospho-STAT3 detection in 96-well format, enabling *in vitro* characterization of thousands of variant combinations in HEK293T cells. In addition, we assessed the effects of selected variants on the four major signaling pathways downstream of LEPR - namely phosphorylation of STAT3, STAT5, ERK1/2, and P70-S6K^22,23^. In sum, these analyses elucidate the functional impact of *LEP* and *LEPR* variants and their interactions, enhancing our understanding of the genetic architecture underlying monogenic forms of obesity and provide functional evidence that may support the existence of combined, possibly digenic, genetic contributions to obesity risk.

## Results

### Assay design and validation for functional characterization of LEP and LEPR variants

Most existing studies characterizing the *LEP/LEPR* pathway rely on Western blotting in HEK293T cells^14,19^. These approaches frequently employ epitope-tagged constructs, yet the potential impact of these tags on protein function is often not systematically assessed. Prior to establishing our functional assay, we therefore evaluated several commonly used tags, including N-terminal Myc, C-terminal Myc, HA, and eYFP-for their influence on leptin receptor signaling (Fig.S1a). Notably, we observed substantial variability in LEPR pathway activation depending on both the type and position of the tag (Fig. S1a). N-terminal Myc tagging preserved or even enhanced signaling, while HA and eYFP tags led to reduced STAT3 phosphorylation compared to the untagged control.

To evaluate whether this tag-dependent effect was cell-line specific, we also performed the assay in HUH7 cells. However, HUH7 cells showed limited dynamic range between LEPR-WT and empty vector controls (Fig. S1b), likely due to high basal STAT3 phosphorylation. Tag effects in HUH7 mirrored those seen in HEK293T, especially with c-Myc increasing and HA/eYFP decreasing signaling. Additionally, the presence of a Myc tag altered the functional classification of certain variants. Specifically, the known pathogenic *LEPR* variant c.1990T>A (p.Trp664Arg) showed apparent rescue with the Myc tag (Fig.1a), potentially leading to a false-negative interpretation of the variant’s impact. As a result, we discontinued the use of tagged *LEPR* constructs and used exclusively untagged wild-type and variant *LEPR* in all subsequent experiments.

For *LEP*, Myc-tagged constructs exhibited higher secretion levels than untagged wild-type leptin (Fig. S1c). However, after normalizing for concentration, the Myc tag did not alter the activity of the wild-type variants (Fig. S1a: 1:50 supernatant of myc-tagged *LEP* reveals the same activity as 1:12 supernatant of untagged *LEP*), consistent with published data ^20,24,25^. Thus, we employed c-Myc–tagged *LEP* to enhance leptin yield in HEK293T cells without compromising the validity of the assay.

To improve throughput and standardization, we designed a custom dot blot device compatible with a 96-well format. We validated this platform against conventional Western blotting using both *LEP* and *LEPR* wild type and known pathogenic variants^19,25–27^. The dot blot assay reliably reproduced differences in signaling intensity observed via Western blotting (Fig. 1b; quantification in Fig. S1d). Both approaches robustly distinguished active signaling from background (Fig. 1c), with only slightly increased variability in dot blot measurements (Fig.S1e).

**Fig. 1:**
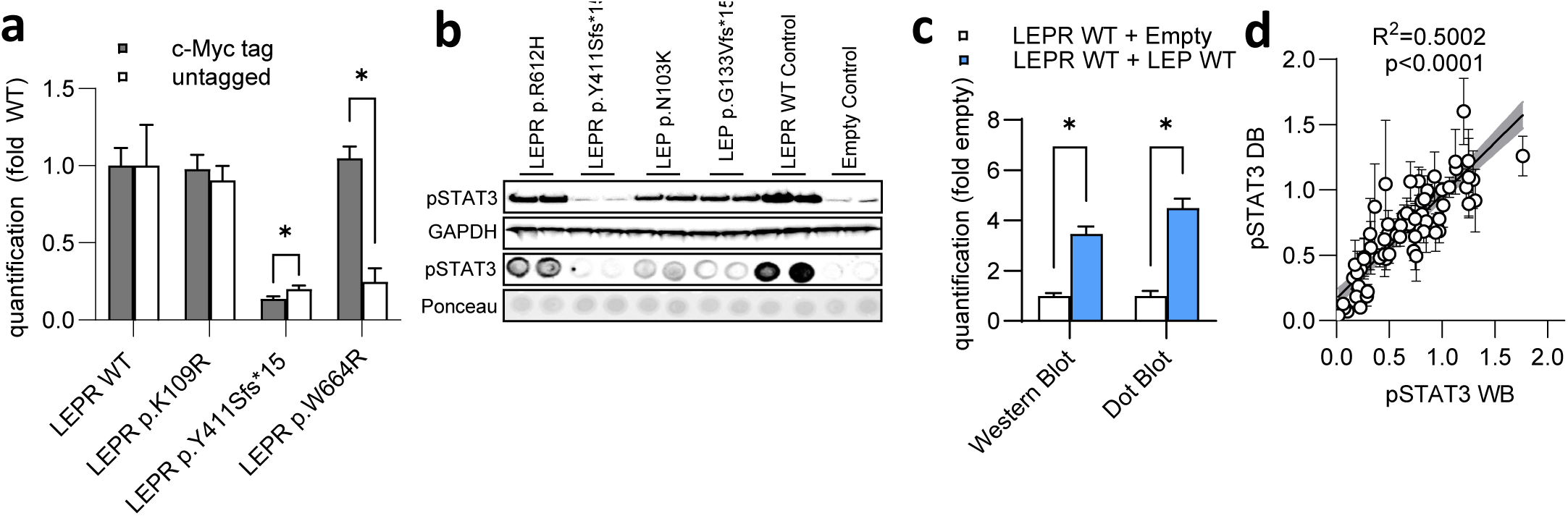
Assay design and validation for functional characterization of *LEP* and *LEPR* variants. (A) Comparison of phosphoStat3 DB data for LEPR WT, p.K109R, p.Y411Sfs*15 and p.W664R both c-terminally myc-tagged and untagged with n=4 in HEK293T cells. (B) Comparison of a Western Blot and a Dot Blot for LEP and LEPR variants. (C) Direct comparison of the DB and WB bands for LEPR WT Control and Empty Control treated with 0.6 ng/ml Leptin Wild Type. (D) Linear regression of our homozygous-like variants measured in Dot Blot and Western Blot.

During the study, we generated side-by-side comparisons of Western blot and dot blot across in sum 65 *LEP* and *LEPR* variants. Quantitative analysis revealed a strong linear correlation between the two methods (Fig. 1d), with comparable but slightly higher standard error in dot blot (Fig.S1e). In sum, both assays were demonstrating analogous feasibility for the use for quantifying ligand and receptor activity. A schematic overview of the assay workflow is provided in Fig. S1f.

### Screening of common *LEP* variants and their compound-heterozygous-like combinations

To evaluate the impact of individual *LEP* variants with the assay, we overexpressed the 30 most common *LEP* variants listed in gnomAD v2.1.1, along with three known pathogenic controls, wild type and empty control, in HEK293T cells (Fig. 2a). Leptin secretion was assessed using ELISA on cell culture supernatants, while intracellular expression was evaluated by anti-leptin Western blotting (Fig. 2b).

**Fig. 2:**
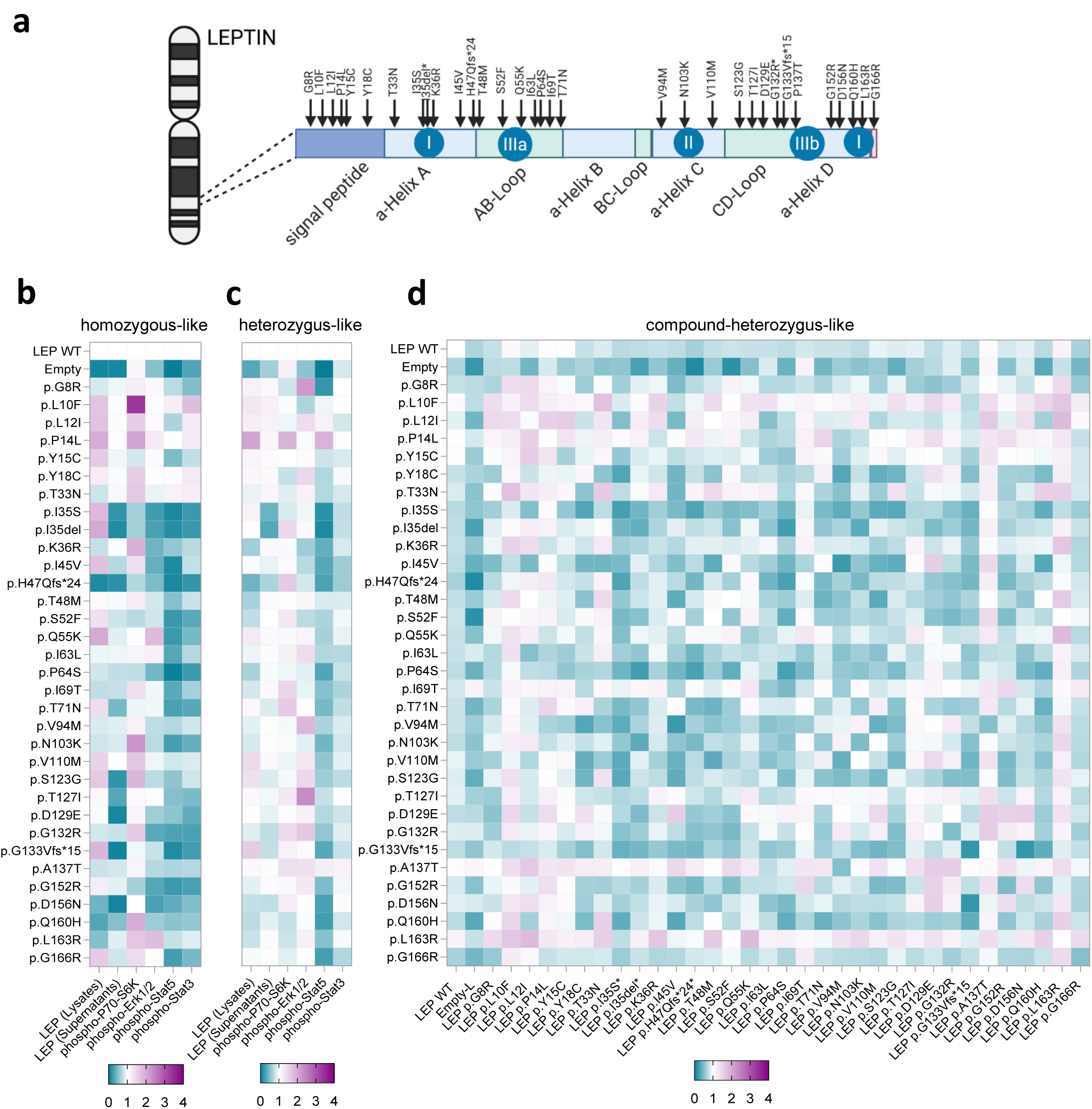
Screening of common leptin variants and their compound-heterozygous-like combinations. (A) Overview of the LEP protein and its domains with the positioning of the researched variants. (B) Treatment of LEPR WT overexpression with supernatants of 35 LEP variants in a homozygous-like mimicking state. Values based on Western Blot (phospho-Proteins, LEP lysates) and ELISA (LEP supernatants) assay and its respective results for Leptin (Lysates), Leptin (Supernatants), phosphoP70-S6K, phosphoErk1/2, phosphoStat5 and phosphoStat3. (C) Treatment of LEPR WT overexpressing cells with supernatants of 35 LEP variants, in a heterozygous-like mimicking state. Values are from Western Blot (phospho-Proteins, LEP lysates) and ELISA (LEP supernatants) assay and its respective results for Leptin (Lysates), Leptin (Supernatants), phosphoP70-S6K, phosphoErk1/2, phosphoStat5 and phosphoStat3. (D) Heatmap of 528 compound-heterozygous-like variant combinations in a phosphoStat3 dot blot

In line with previous reports, pathogenic variants like p.Gly133ValfsTer15^16^, p.Ile35Ser^25^ and p.Ile35del showed absent secretion^20^, whereas other reported pathogenic variants^20^, such as p.Pro64Ser (MAF=0%) and p.Asn103Lys^27^ (MAF=0.004151%) revealed preserved secretion, indicating a defect in a downstream mechanism.

From the other investigated variants, one variant, p.His47GlnfsTer24 (gnomAD v4.1.0 MAF=0.00006196%), was undetectable in both supernatant and lysate. Two variants, p.Thr71Asn (MAF=0.0003098%) and p.Thr127Ile (MAF=0.0003098%), showed markedly reduced secretion. Another variant, p.Gln160His (MAF=0.0004343%), exhibited low intracellular expression but retained partial secretion. A larger subset of variants - including p.Ile35Ser (MAF=0%), p.Ile35del (MAF=0.0001239%), p.Ser123Gly (MAF=0.0001239%), p.Asp129Glu (MAF=0.001921%), p.Gly133ValfsTer15 (MAF=0.0008055%), and p.Asp156Asn (MAF=0.0008680%) - were detectable intracellularly but absent in the secretion assay.

Collectively, these findings support the pathogenic potential of several previously uncharacterized variants - p.His47GlnfsTer24, p.Thr71Asn, p.Ser123Gly, p.Thr127Ile, p.Asp129Glu, and p.Asp156Asn - that may result in classical leptin deficiency when present in homozygous form.

To mimic a heterozygous-like condition, we co-expressed each *LEP* variant with WT *LEP*. In most cases, except for p.Ile35Ser, p.Ile35del, and p.His47GlnfsTer24, leptin secretion was restored to near-normal levels, consistent with clinical observations that biallelic pathogenic variants are required to manifest the full leptin deficiency phenotype (Fig. 2c).

To assess biological activity, we treated HEK293T cells overexpressing WT *LEPR* with variant-containing supernatants. Variants with impaired secretion led to a marked loss of signaling via phosphorylated STAT3, STAT5, and ERK1/2, while phosphorylation of P70-S6K remained largely unaffected. Additionally, several secreted *LEP* variants - p.Gly8Arg (MAF=0.002230%), p.Ser52Phe (MAF=0.0001240%), p.Gln55Lys (MAF=0.0006197%), p.Pro64Ser, p.Asn103Lys, p.Gly132Arg (MAF=0.0008054%), p.Gly152Arg (MAF=0.0002479%), and p.Gly166Arg (MAF=0.009196%) - demonstrated significant reductions in STAT3 phosphorylation under homozygous-like expression conditions (Fig. 2b).

Given that most variants are likely inherited in heterozygous form in the population, we also assessed coexpressed WT/variant supernatants. As expected, WT co-expression improved signaling in most cases. Interestingly, phospho-STAT5 activity remained reduced (Fig. 2c).

We defined the pathogenicity of the variant activity as being lower than that of published pathogenic variants, which all displayed activity below 50% (Fig.S2a). Given this threshold, a total of 12 previously uncharacterized *LEP* variants showed significantly reduced STAT3 phosphorylation (p < 0.05), each displaying <50% of WT activity (Fig. S2a). Functional findings for p.Ile35Ser, p.Ile35del, p.Pro64Ser, p.Asn103Lys, and p.Gly133ValfsTer15 aligned well with published data^20^, underscoring the robustness and reproducibility of our assay in comparison to existing clinical and experimental literature.

We next evaluated all 528 possible compound-heterozygous-like combinations of the tested *LEP* variants using dot blot (Fig. 2d). Combinations with notable effects were re-tested with increased replicates for validation. In total, 57 combinations - predominantly involving known or suspected loss-of-function alleles - resulted in a marked loss of STAT3 activation.

Interestingly, several combinations involving p.Val94Met (MAF=0.4662%) or p.Val110Met (MAF=0.01524%) - both common variants with minimal effect in homozygous-like states (Fig. 2b) - nearly exhibited abolished signaling when paired with other mildly hypomorphic variants (Fig. 2d). These findings suggest that such variants may act as functional risk alleles in compound-heterozygous state, despite appearing benign individually.

### Screening of common leptin-receptor variants and their compound-heterozygous-like combination

To evaluate the functional impact of common *LEPR* variants, we overexpressed the 19 most frequent alleles, 3 known-pathogenic control variants, 3 further frameshift variants to represent this large group of individually quite rare variants, 6 intracellular phosphorylation site variants, wild type and empty control (Fig. 3a) in HEK293T cells and stimulated them with wild-type leptin-containing supernatants. As expected, the known pathogenic variants p.Tyr411SerfsTer15^19^ (MAF=0%), p.Arg612His^28^ (MAF=0.03872%), and p.Trp664Arg^26^ (MAF=0.001056%) exhibited markedly reduced phosphorylation of STAT3, STAT5, and ERK1/2, consistent with previous reports. In contrast, P70-S6K phosphorylation was variably affected (Fig. 3b).

**Fig. 3:**
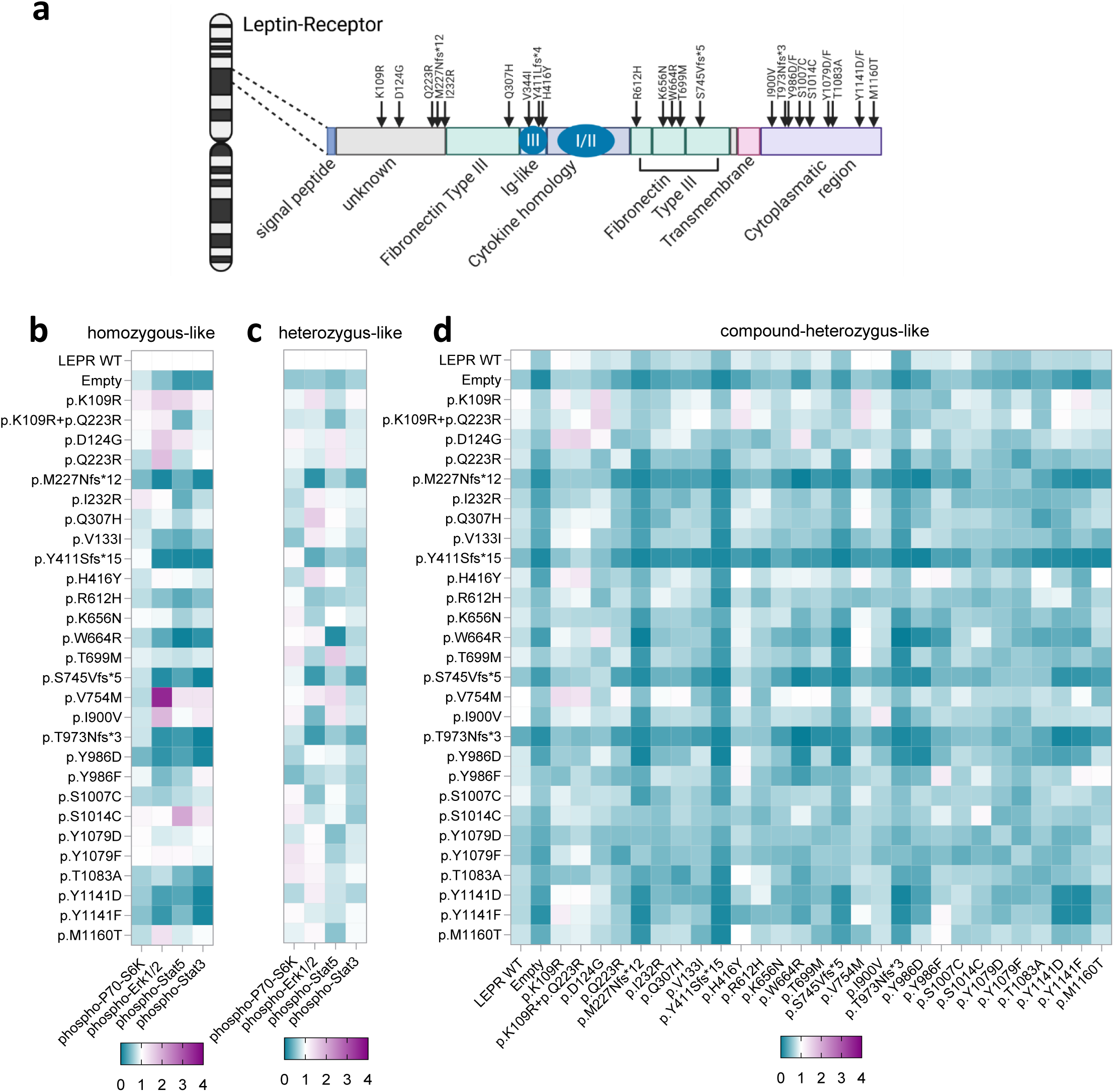
Screening of common leptin-receptor variants and their compound-heterozygous-like combinations. (A) Overview of the LEPR protein and its domains with the positioning of the researched variants. (B) Overexpression of 30 LEPR variants in a homozygous-like state. Values are derived from Western Blot and its respective results for phosphoP70-S6K, phosphoErk1/2, phosphoStat5 and phosphoStat3 are indicated. (C) Co-expression of 30 LEPR variants in a heterozygous-like in-vitro assay and its respective results for phosphoP70-S6K, phosphoErk1/2, phosphoStat5 and phosphoStat3. (D) Heatmap of 406 compound-heterozygous-like variant combinations in a phosphoStat3 DB.

Among the tested variants, seven - p.Met227AsnfsTer12 (MAF=0.0006819%), p.Ser745ValfsTer4 (MAF=0.0006829%), p.Thr973AsnfsTer3 (MAF=0.00006195%), p.Tyr986Asp (MAF=0%), p.Thr1083Ala (MAF=0.01363%), p.Tyr1141Asp (MAF=0%), and p.Tyr1141Phe (MAF=0%) - demonstrated consistently and severely impaired signaling activity (Fig. 3b). Notably, two variants, p.Val754Met (MAF=0.1013%) and p.Ile900Val (MAF=0.05484%), showed increased activity, particularly through enhanced ERK1/2 phosphorylation (Fig. 3b). Given the absence of reported gain-of-function phenotypes for *LEPR*, the (patho-) physiological relevance of these variants remains to be further investigated.

In line with prior studies^29^, co-expression of WT *LEPR* with the respective missense variants restored signaling across all tested pathways (Fig. 3b), although some clinical data suggest partial phenotypic effects in heterozygous carriers^17^. In contrast, frameshift variants were not fully rescued, highlighting a functional distinction between missense and frameshift mutations in *LEPR*.

Additionally, we identified seven previously uncharacterized *LEPR* variants with significantly reduced function (p < 0.05), each exhibiting less than 50% of WT STAT3 phosphorylation activity when tested in a homozygous-like context (Fig.S2b).

A screen of 406 compound-heterozygous-like *LEPR* variant combinations identified 145 pairs falling within the pathogenic range, as defined by the STAT3 phosphorylation threshold established for homozygous-like expression of known pathogenic variants (Fig. 3d). The majority of these combinations involved one of the four frameshift variants tested, although certain missense–missense pairs, which were previously shown to impair signaling individually, also contributed to the pathogenic combination pool. Notably, several combinations of extracellular (e.g., p.Arg612His, p.Trp664Arg) and intracellular (e.g., p.Tyr986Asp) missense variants exhibited higher STAT3 activity than either variant alone, indicating potential partial functional complementation between distinct *LEPR* domains.

### Digenic interactions between *LEP* and *LEPR* variants reveal extensive functional impairment

To explore potential digenic effects, we evaluated 1,050 homozygous-like (Fig. 4a) or heterozygous-like (Fig. 4b) variant combinations involving one *LEP* and one *LEPR* variant. Here we expressed either the *LEP* or *LEPR* variants alone (Fig. 4a), or each co-expressed with its corresponding wild type allele to mimic the heterozygous state (Fig. 4b).

**Fig. 4:**
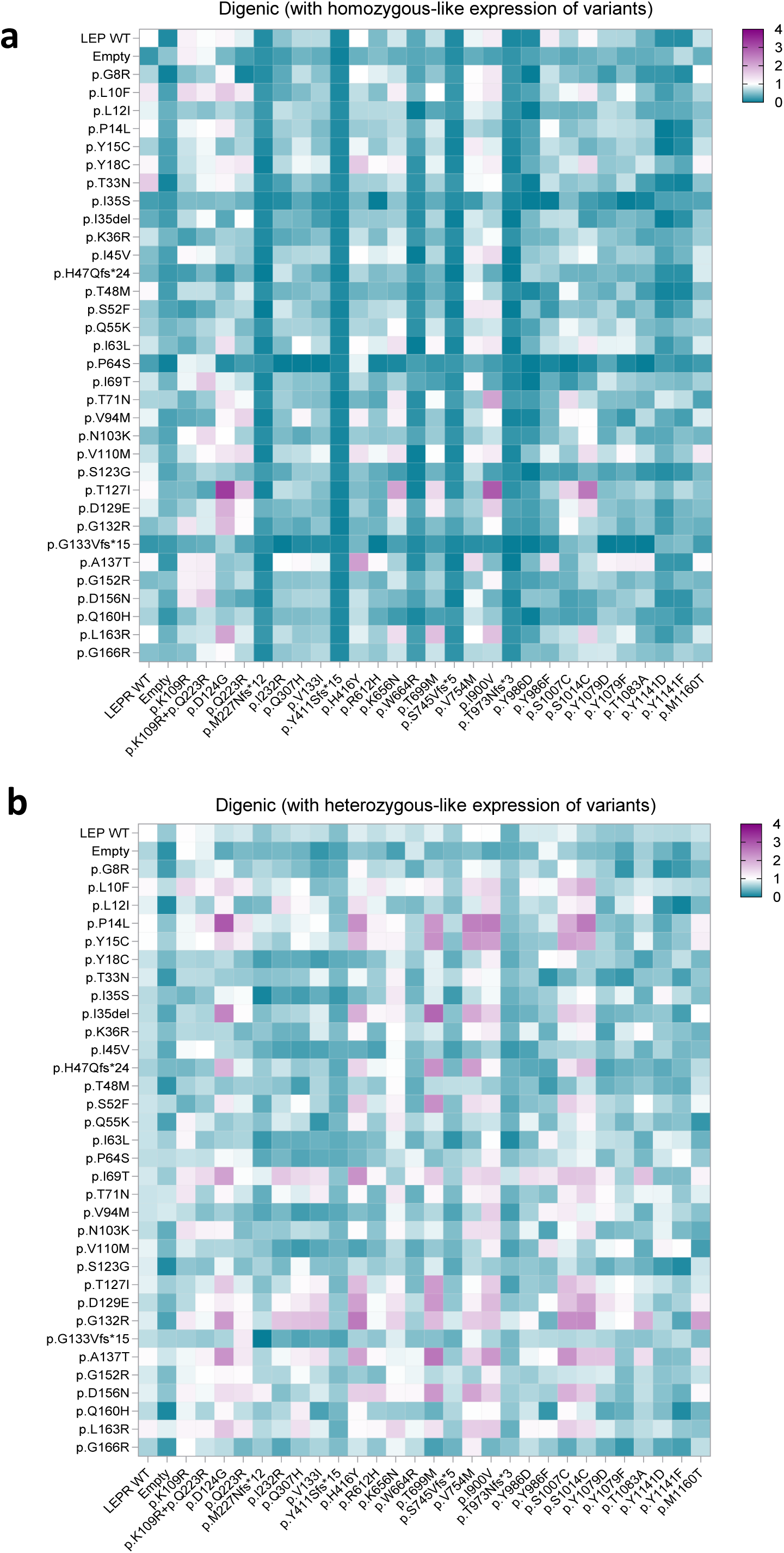
Digenic interactions between *LEP* and *LEPR* variants reveal extensive functional impairment. (A) Heatmap of 1049 digenic homozygous-like variant combinations in a phosphoStat3 DB. (B) Heatmap of 1050 digenic heterozygous-like variant combinations in a phosphoStat3 DB.

Although genotypes of homozygous-like digenic combinations are not described in clinical settings, they provide valuable mechanistic insight. As expected, combinations involving fully nonfunctional *LEP* and *LEPR* variants resulted in complete loss of STAT3 signaling, which was not rescued by co-expression with either a wild type allele of the second gene or a putative gain-of-function variant in the second gene. In total, 534 of the 1,050 digenic homozygous-like combinations fell below the pathogenic cut-off, underscoring the strict interdependence of *LEP* and *LEPR* for leptin receptor activation.

When both gene variants in *LEP* and *LEPR* were coexpressed with a wild type allele, which is a combination with higher likelihood to be found in clinical settings, still 272 combinations exhibited STAT3 phosphorylation below the predefined pathogenic threshold of <50% WT activity. As anticipated after seeing their significant STAT3 hypophosphorylation in a heterozygous-like state in western blot and in a compound-heterozygous-like state in dot blot, *LEPR* frameshift variants were overrepresented among dysfunctional combinations. Interestingly, several pairings involving *LEPR* variants with p.Val754Met or p.Ile900Val, or *LEP* variants with p.Pro14Leu (MAP=0.0001239%) or p.Leu10Phe (MAP=0.0008674%), resulted in consistently elevated STAT3 phosphorylation, suggesting a potential gain-of-function effect.

### *In silico* predictors show poor correlation with *LEP* and *LEPR* functional phenotypes

To evaluate the performance of bioinformatic tools in predicting functional outcomes, we compared our experimental phospho-STAT3 data from Western blotting with commonly used *in silico* prediction scores - including CADD, REVEL, and PolyPhen-2 - for all missense and frameshift variants available in our dataset.

Linear regression analyses (Fig. 5a-c) revealed the absence of significant association between STAT3 phosphorylation and any of the three *in silico* scores. These findings challenge the current utility of widely used computational predictors in assessing *LEP* and *LEPR* variants, particularly in relation to STAT3 activity, the primary mediator of Leptin receptor signaling, due to its role in initiating transcription of both *POMC* and *SOCS3* in the hypothalamic arcuate nucleus^30–32^.

**Fig. 5:**
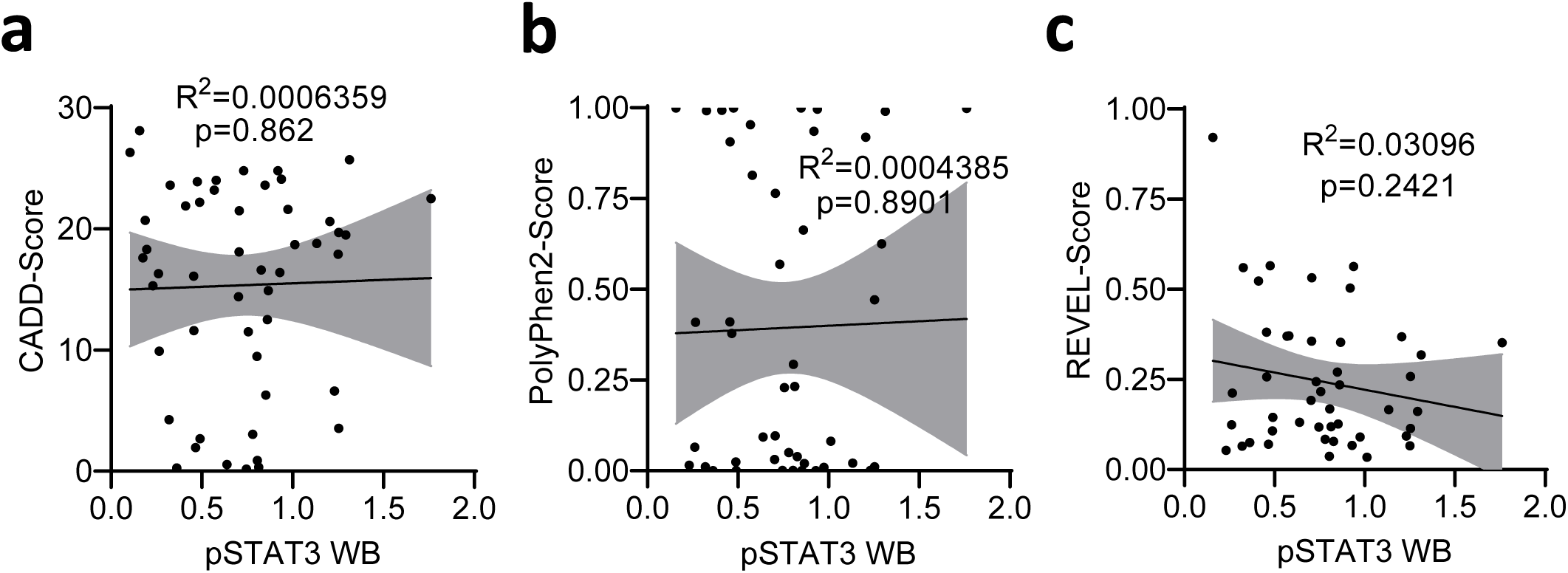
Comparison to *in silico* prediction tools. (A) Linear regression of CADD-Score and phosphoStat3 WB data. (B) Linear regression of PolyPhen-2-Score and phosphoStat3 WB data. (C) Linear regression of REVEL-Score and phosphoStat3 WB data.

Taken together, our results highlight the limitations of current *in silico* tools in the context of hormonal signaling genes and underscore the need for improved algorithms specifically calibrated for functionally constrained pathways such as *LEP/LEPR* and the use of biological variant characterization.

## Discussion

Our study establishes a simple, scalable assay for functional assessment of *LEP* and *LEPR* variants, enabling systematic analysis of VUS in the heterozygous, biallelic, and digenic context. This approach addresses a critical gap in functional analysis of obesity-related genetic diagnostics, where numerous VUS confound clinical interpretation^33^. By directly quantifying receptor signaling via STAT3 phosphorylation, our platform provides robust measurements to classify variants comparable to established assays and beyond prediction tools.

Applying this assay, we identified 19 previously uncharacterized hypofunctional variants in *LEP* and *LEPR* and provided a comprehensive map of functional interactions between variants in homozygous-like, compound-heterozygous-like, and digenic contexts. Here, variants, e.g. *LEP* p.Val110Met or p.Val94Met, can exacerbate their dysfunction when paired with other dysfunctional alleles, unmasking their potentially pathogenic effects only in the context of other variants - which has been described for other genes^34–36^. Moreover, we found that digenic *LEP/LEPR* combinations often reduced signaling below levels seen for known pathogenic variants, supporting emerging evidence that multiple heterozygous variants in the leptin–melanocortin pathway elevate obesity risk^37,38^. Of note, we also identified potential gain-of-function variants that could be explored in the context of thinness and anorexia in future studies^39^.

Our results also underscore the significant impact of protein tagging on LEPR signaling. We observed that tags can introduce artifacts, leading to both under- and overestimation of variant pathogenicity, which highlights the need for validation using untagged constructs. Discrepancies between the use of tagged and untagged constructs may explain conflicting results across studies and emphasize the importance of physiologically relevant assay design. This is especially important in the context of discordant findings on the STAT3 phosphorylation levels of c.1231_1233 (p.Tyr411Serfs*15) with constructs tagged with myc-tag and eYFP^19^. Additional studies used mostly n-terminally myc-tagged *LEPR* constructs^28,40^ but also other tags such as FLAG, c-terminal myc^41^ and YFP derivates^42^.

Nonetheless, our study has limitations that may affect the direct translation of our findings to *in vivo* contexts. We exclusively expressed the LEPRb isoform, which is found on several cell types including the Nucleus arcuatus neurons^43,44^. However, shorter isoforms have been described^42^. Since they lack a great portion of the LEPRb signalling properties^45^, it unmasks their use as an artificial overexpression system. In the same line, we conducted our experiments in plasmid-transfected HEK293T cells rather than in neurons, so certain cell-type- or translation-specific effects may not have been captured.

In summary, we expand the catalog of pathogenic *LEP* and *LEPR* variants and define multiple mechanisms of dysfunction, including hyposecretion, hypofunction, and domain-specific complementation in heterozygous, biallelic and digenic contexts. Further validation in population-based or clinical cohorts will be important to determine the clinical relevance of these findings.

Ultimately, integrating this scalable functional assay into diagnostic workflows could advance personalized medicine by enabling rapid assessment of uncharacterized variants, variant combinations, and digenic interactions in patients undergoing genetic testing for hormone or receptor-associated diseases.

## Materials and Methods

### Variant Inclusion Criteria

All variants were selected based on stringent criteria to maximize diagnostic applicability. Wild-type reference sequences for *LEPR* (NM_002303.6) and *LEP* (NM_000230.3), the respective empty control vector, and three previously validated pathogenic controls for each gene were included. In addition, the 19 most common coding variants in *LEPR* and 30 in *LEP*, as reported in gnomAD v.2.1.1, were analyzed. To represent the cumulative prevalence of *LEPR* frameshift variants, three additional frameshift alleles - distributed across the receptor - were included. Six phosphorylation site variants were incorporated to capture intracellular functional effects.

### Cell Culture

HEK293T and HuH-7 cells were cultured in high-glucose DMEM supplemented with 10% fetal calf serum (Sigma) and 1% antibiotic-antimycotic solution (Sigma). Transfections were performed using jetOptimus® reagent (polyplus). Four hours post-transfection, the medium was refreshed. Experiments were conducted the following day at 60–80% confluency.

### Plasmids

All purchased constructs (listed below) were designed and cloned into a pcDNA3.1(+) backbone, with or without added tag (GenScript).

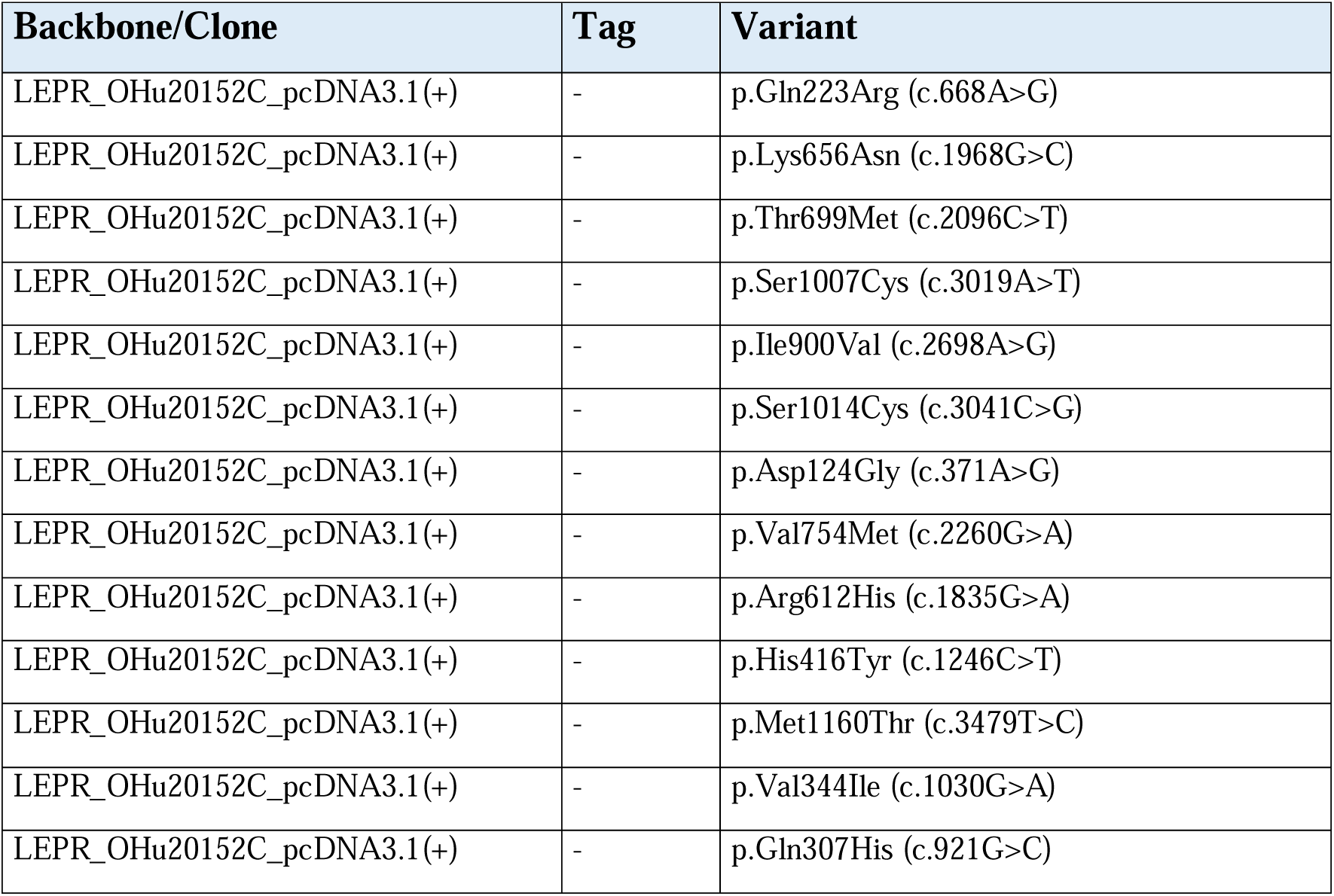
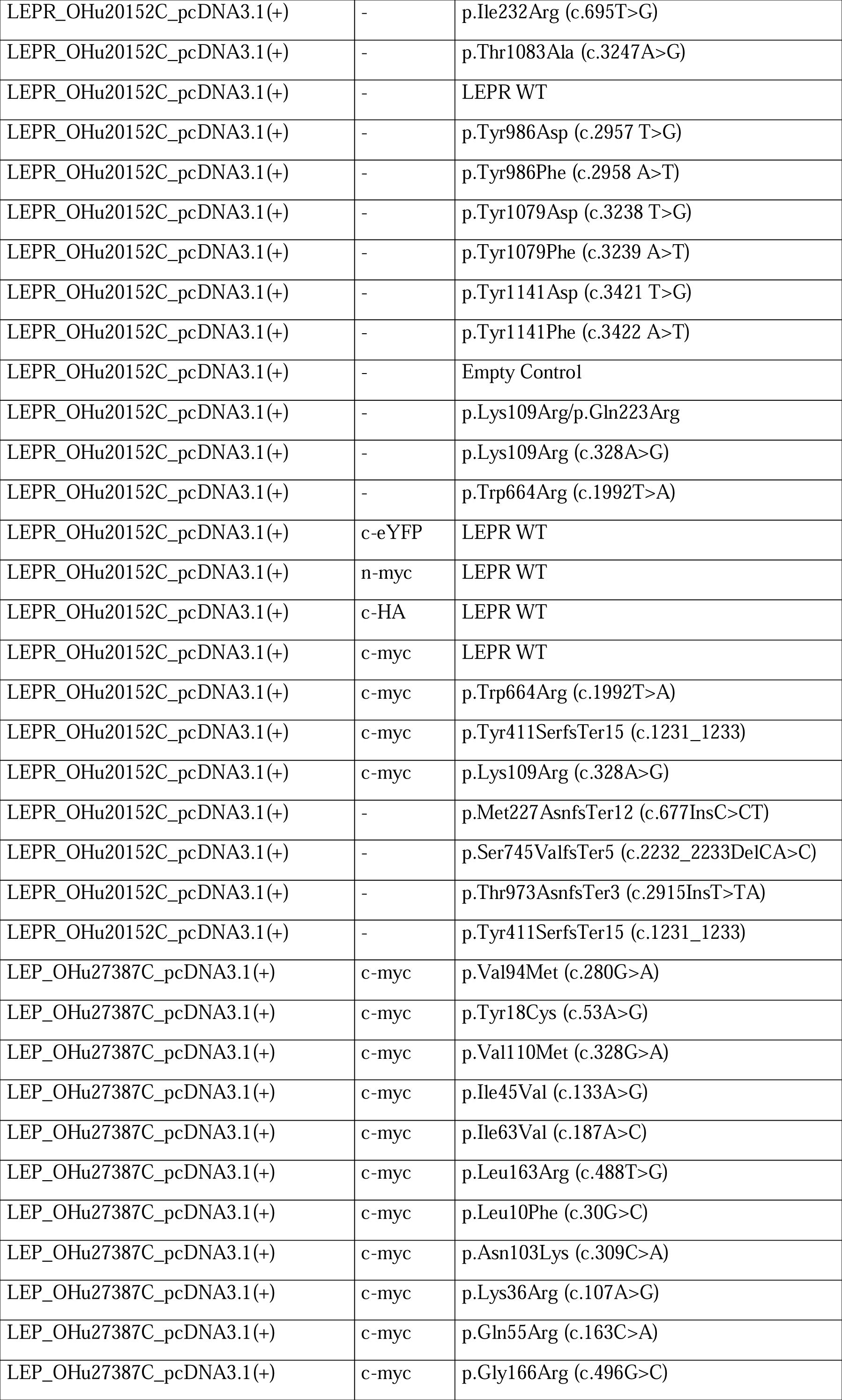
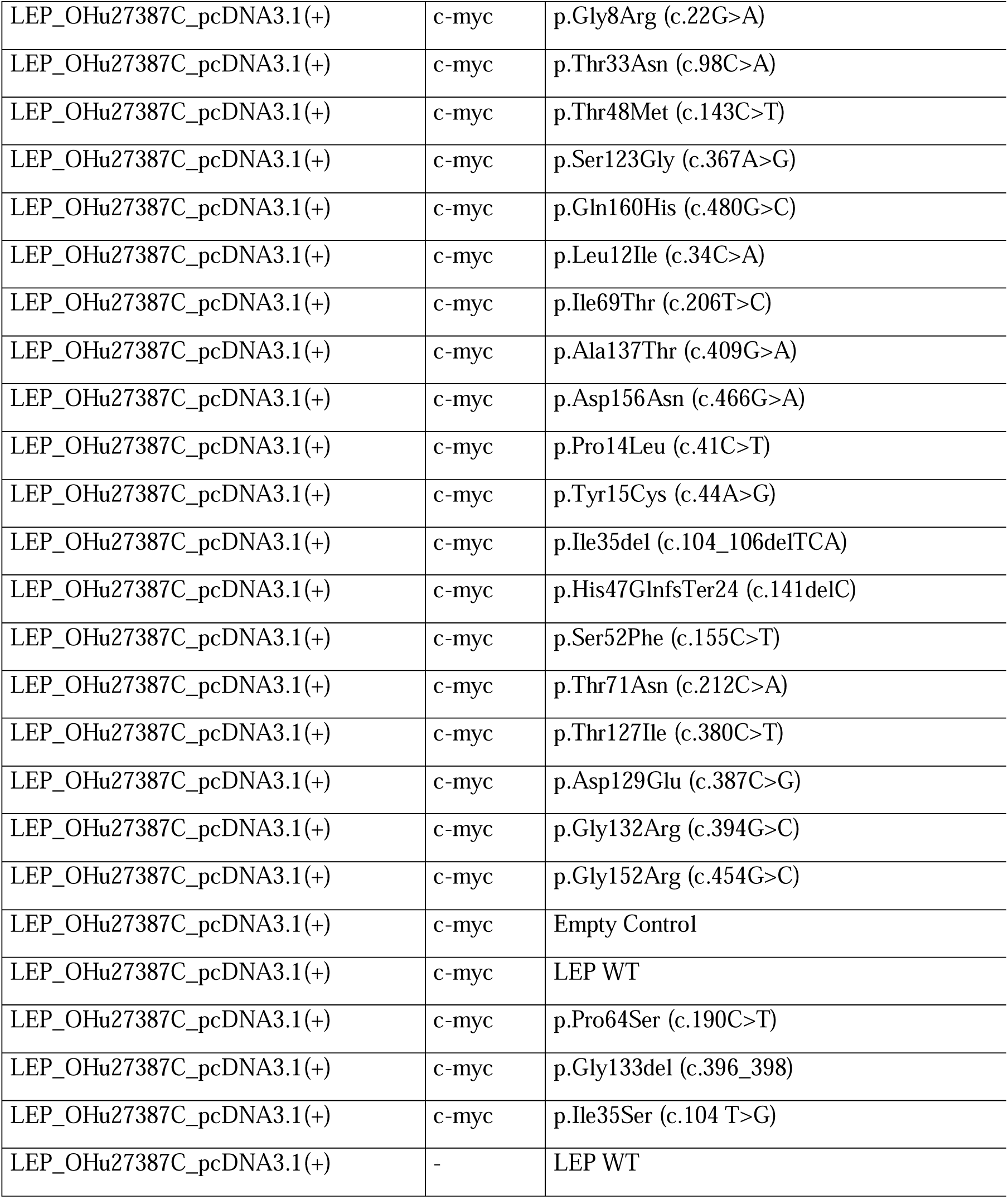

| Backbone/Clone | Tag | Variant |
| --- | --- | --- |
| LEPR_OHu20152C_pcDNA3.1(+) | - | p.Gln223Arg (c.668A>G) |
| LEPR_OHu20152C_pcDNA3.1(+) | - | p.Lys656Asn (c.1968G>C) |
| LEPR_OHu20152C_pcDNA3.1(+) | - | p.Thr699Met (c.2096C>T) |
| LEPR_OHu20152C_pcDNA3.1(+) | - | p.Ser1007Cys (c.3019A>T) |
| LEPR_OHu20152C_pcDNA3.1(+) | - | p.Ile900Val (c.2698A>G) |
| LEPR_OHu20152C_pcDNA3.1(+) | - | p.Ser1014Cys (c.3041C>G) |
| LEPR_OHu20152C_pcDNA3.1(+) | - | p.Asp124Gly (c.371A>G) |
| LEPR_OHu20152C_pcDNA3.1(+) | - | p.Val754Met (c.2260G>A) |
| LEPR_OHu20152C_pcDNA3.1(+) | - | p.Arg612His (c.1835G>A) |
| LEPR_OHu20152C_pcDNA3.1(+) | - | p.His416Tyr (c.1246C>T) |
| LEPR_OHu20152C_pcDNA3.1(+) | - | p.Met1160Thr (c.3479T>C) |
| LEPR_OHu20152C_pcDNA3.1(+) | - | p.Val344Ile (c.1030G>A) |
| LEPR_OHu20152C_pcDNA3.1(+) | - | p.Gln307His (c.921G>C) |
| LEPR_OHu20152C_pcDNA3.1(+) | - | p.Ile232Arg (c.695T>G) |
| LEPR_OHu20152C_pcDNA3.1(+) | - | p.Thr1083Ala (c.3247A>G) |
| LEPR_OHu20152C_pcDNA3.1(+) | - | LEPR WT |
| LEPR_OHu20152C_pcDNA3.1(+) | - | p.Tyr986Asp (c.2957 T>G) |
| LEPR_OHu20152C_pcDNA3.1(+) | - | p.Tyr986Phe (c.2958 A>T) |
| LEPR_OHu20152C_pcDNA3.1(+) | - | p.Tyr1079Asp (c.3238 T>G) |
| LEPR_OHu20152C_pcDNA3.1(+) | - | p.Tyr1079Phe (c.3239 A>T) |
| LEPR_OHu20152C_pcDNA3.1(+) | - | p.Tyr1141Asp (c.3421 T>G) |
| LEPR_OHu20152C_pcDNA3.1(+) | - | p.Tyr1141Phe (c.3422 A>T) |
| LEPR_OHu20152C_pcDNA3.1(+) | - | Empty Control |
| LEPR_OHu20152C_pcDNA3.1(+) | - | p.Lys109Arg/p.Gln223Arg |
| LEPR_OHu20152C_pcDNA3.1(+) | - | p.Lys109Arg (c.328A>G) |
| LEPR_OHu20152C_pcDNA3.1(+) | - | p.Trp664Arg (c.1992T>A) |
| LEPR_OHu20152C_pcDNA3.1(+) | c-eYFP | LEPR WT |
| LEPR_OHu20152C_pcDNA3.1(+) | n-myc | LEPR WT |
| LEPR_OHu20152C_pcDNA3.1(+) | c-HA | LEPR WT |
| LEPR_OHu20152C_pcDNA3.1(+) | c-myc | LEPR WT |
| LEPR_OHu20152C_pcDNA3.1(+) | c-myc | p.Trp664Arg (c.1992T>A) |
| LEPR_OHu20152C_pcDNA3.1(+) | c-myc | p.Tyr411SerfsTer15 (c.1231_1233) |
| LEPR_OHu20152C_pcDNA3.1(+) | c-myc | p.Lys109Arg (c.328A>G) |
| LEPR_OHu20152C_pcDNA3.1(+) | - | p.Met227AsnfsTer12 (c.677InsC>CT) |
| LEPR_OHu20152C_pcDNA3.1(+) | - | p.Ser745ValfsTer5 (c.2232_2233DelCA>C) |
| LEPR_OHu20152C_pcDNA3.1(+) | - | p.Thr973AsnfsTer3 (c.2915InsT>TA) |
| LEPR_OHu20152C_pcDNA3.1(+) | - | p.Tyr411SerfsTer15 (c.1231_1233) |
| LEP_OHu27387C_pcDNA3.1(+) | c-myc | p.Val94Met (c.280G>A) |
| LEP_OHu27387C_pcDNA3.1(+) | c-myc | p.Tyr18Cys (c.53A>G) |
| LEP_OHu27387C_pcDNA3.1(+) | c-myc | p.Val110Met (c.328G>A) |
| LEP_OHu27387C_pcDNA3.1(+) | c-myc | p.Ile45Val (c.133A>G) |
| LEP_OHu27387C_pcDNA3.1(+) | c-myc | p.Ile63Val (c.187A>C) |
| LEP_OHu27387C_pcDNA3.1(+) | c-myc | p.Leu163Arg (c.488T>G) |
| LEP_OHu27387C_pcDNA3.1(+) | c-myc | p.Leu10Phe (c.30G>C) |
| LEP_OHu27387C_pcDNA3.1(+) | c-myc | p.Asn103Lys (c.309C>A) |
| LEP_OHu27387C_pcDNA3.1(+) | c-myc | p.Lys36Arg (c.107A>G) |
| LEP_OHu27387C_pcDNA3.1(+) | c-myc | p.Gln55Arg (c.163C>A) |
| LEP_OHu27387C_pcDNA3.1(+) | c-myc | p.Gly166Arg (c.496G>C) |
| LEP_OHu27387C_pcDNA3.1(+) | c-myc | p.Gly8Arg (c.22G>A) |
| LEP_OHu27387C_pcDNA3.1(+) | c-myc | p.Thr33Asn (c.98C>A) |
| LEP_OHu27387C_pcDNA3.1(+) | c-myc | p.Thr48Met (c.143C>T) |
| LEP_OHu27387C_pcDNA3.1(+) | c-myc | p.Ser123Gly (c.367A>G) |
| LEP_OHu27387C_pcDNA3.1(+) | c-myc | p.Gln160His (c.480G>C) |
| LEP_OHu27387C_pcDNA3.1(+) | c-myc | p.Leu12Ile (c.34C>A) |
| LEP_OHu27387C_pcDNA3.1(+) | c-myc | p.Ile69Thr (c.206T>C) |
| LEP_OHu27387C_pcDNA3.1(+) | c-myc | p.Ala137Thr (c.409G>A) |
| LEP_OHu27387C_pcDNA3.1(+) | c-myc | p.Asp156Asn (c.466G>A) |
| LEP_OHu27387C_pcDNA3.1(+) | c-myc | p.Pro14Leu (c.41C>T) |
| LEP_OHu27387C_pcDNA3.1(+) | c-myc | p.Tyr15Cys (c.44A>G) |
| LEP_OHu27387C_pcDNA3.1(+) | c-myc | p.Ile35del (c.104_106delTCA) |
| LEP_OHu27387C_pcDNA3.1(+) | c-myc | p.His47GlnfsTer24 (c.141delC) |
| LEP_OHu27387C_pcDNA3.1(+) | c-myc | p.Ser52Phe (c.155C>T) |
| LEP_OHu27387C_pcDNA3.1(+) | c-myc | p.Thr71Asn (c.212C>A) |
| LEP_OHu27387C_pcDNA3.1(+) | c-myc | p.Thr127Ile (c.380C>T) |
| LEP_OHu27387C_pcDNA3.1(+) | c-myc | p.Asp129Glu (c.387C>G) |
| LEP_OHu27387C_pcDNA3.1(+) | c-myc | p.Gly132Arg (c.394G>C) |
| LEP_OHu27387C_pcDNA3.1(+) | c-myc | p.Gly152Arg (c.454G>C) |
| LEP_OHu27387C_pcDNA3.1(+) | c-myc | Empty Control |
| LEP_OHu27387C_pcDNA3.1(+) | c-myc | LEP WT |
| LEP_OHu27387C_pcDNA3.1(+) | c-myc | p.Pro64Ser (c.190C>T) |
| LEP_OHu27387C_pcDNA3.1(+) | c-myc | p.Gly133del (c.396_398) |
| LEP_OHu27387C_pcDNA3.1(+) | c-myc | p.Ile35Ser (c.104 T>G) |
| LEP_OHu27387C_pcDNA3.1(+) | - | LEP WT |

### Leptin Supernatants

Supernatants were generated from HEK293T cells seeded at 100,000 cells per well in 48-well plates. Each variant was replicated across four independent plates. After transfection and medium change, cells were incubated for 20 hours before supernatant collection. Samples were diluted 1:1 and checked by Leptin-ELISA, and Western blot agains c-Myc and Leptin. For functional testing in signaling assays, we used this diluted and pooled supernatant at a 1:50 working dilution in the cell culture media, corresponding to 0.6 ng/mL wild-type leptin, to reflect physiological leptin concentrations in cerebrospinal fluid^46,47^.

### Western Blot

Cells were cultured as specified and treated with Leptin supernatants for 45 minutes prior to cell lysis. After washing with PBS, cells were directly lysed in 4× Laemmli buffer (30% glycerol, 0.3M DTT, 6.7% SDS, 0.01% bromophenol blue, 80 nM Tris-HCl pH 6.8, plus complete Mini Protease Inhibitors and Phosphatase Inhibitors (Roche)) and incubated for 20 minutes on wet ice. Centrifugtion for 5 minutes at 4°C at 2000xG followed before they were transferred to sample tubes. Leptin supernatants were supplemented with 1:4 4x Laemmli Buffer. Both lysates and supernatants were then boiled at 95°C for 5 minutes and then processed by protein separation on SDS-polyacrylamide gels and transfer to PVDF membranes using the Transblot Turbo Transfer System (Bio-Rad laboratories). Following blocking (20 mM Tris-HCl, pH 7.4; 150 mM NaCl; 0.1% Tween-20; 5% non-fat dry milk) and washing (20 mM Tris–HCl, pH 7.4; 150 mM NaCl; 0.1% Tween-20), the membranes were incubated in primary antibody solution (20 mM Tris–HCl, pH 7.4; 150 mM NaCl; 0.1% Tween-20; 5% BSA or 2.5% non-fat dry milk) containing the respective antibodies overnight. Membranes were washed again in TBS-T and incubated with appropriate secondary HRP-congjugated antibodies for an hour. After final washing, proteins were visualized using the ChemiDoc MP Imaging System (Bio-Rad laboratories) and quantified with ImageJ. Sample data was subtracted from background levels and normalized to wild-type controls. The following antibodies were used:

| Target | Company | Catalogue Number | Dilution Western Blot | Dilution Dot Blot |
| --- | --- | --- | --- | --- |
| phospho-P70-S6K | Cell Signaling | #9205 | 1:1000 | 1:800 |
| P70-S6K (Total) | Cell Signaling | #9202 | 1:1000 | - |
| phospho-Erk1/2 | Cell Signaling | #9101 | 1:1000 | - |
| Erk1/2 (Total) | Cell Signaling | #9102 | 1:1000 | 1:500 |
| phospho-Stat5 | Cell Signaling | C11C6 | 1:500 | - |
| Stat5 (Total) | Cell Signaling | D2O6Y | 1:1000 | - |
| phospho-Stat3 | Cell Signaling | M9C6 | 1:1000 | 1:800 |
| Stat3 (Total) | Cell Signaling | 79D7 | 1:1000 | - |
| Gapdh | abcam | ab8245 | 1:1000 | 1:800 |
| Leptin | antibodies-online Inc. | ABIN7445093 | 1:200 | - |
| myc-tag | Merck | M5549 | 1:100 | - |
| rabbit IgG | Cell Signaling | #7074 | 1:2000 | 1:1000 |
| mouse IgG | Cell Signaling | #7076 | 1:2000 | 1:1000 |

### Dot Blot

Cells were cultured as specified and treated with Leptin supernatants for 45 minutes prior to cell lysis. After washing with PBS, cells were directly lysed in 4× Laemmli buffer (30% glycerol, 0.3M DTT, 6.7% SDS, 0.01% bromophenol blue, 80 nM Tris-HCl pH 6.8, plus complete Mini Protease Inhibitors and Phosphatase Inhibitors (Roche)) and incubated for 20 minutes on wet ice. Centrifugtion for 5 minutes at 4°C at 2000xG followed before they were transferred to 96-Well-PCR-Sample Plates to be boiled for 15 minutes in a PCR Cycler at 95°C. Boiled lysates were applied to a nitrocellulose membrane (Amersham™, Cytiva) via a custom 3D-printed 96-well slot blot device (BambuLab X1C) at 6 μl per dot. Following blocking (20 mM Tris–HCl, pH 7.4; 150 mM NaCl; 0.1% Tween-20; 5% non-fat dry milk), washing (20 mM Tris–HCl, pH 7.4; 150 mM NaCl; 0.1% Tween-20) and shaking in WesternBright ECL HRP substrate, membranes were incubated in primary antibody solution (phospho-Stat3 (M9C6, Cell Signaling Technology®) 1:800 in 20 mM Tris–HCl, pH 7.4; 150 mM NaCl; 0.1% Tween-20; 5% BSA) overnight. Following washing, cells were incubated with the appropriate HRP-conjugated secondary antibody 1:1000 in Blocking Buffer. After final washing, proteins were visualized using the ChemiDoc MP Imaging System (Bio-Rad laboratories) and quantified with ImageJ. Sample data was subtracted from background levels and normalized to wild-type controls.

### Statistical Methods

Statistical analyses were performed with GraphPad Prism v10.1.2. Comparisons to wild-type controls were conducted using unpaired t-tests with Welch correction, followed by correction for multiple comparisons using the Benjamini–Krieger–Yekutieli test. Bars are presented as mean ± standard error of the mean (SEM). Correlations were assessed using linear regression, with reporting of confidence intervals, p-values, and R² values.

## Acknowledgement

The authors thank Katrin Rading for excellent technical support. C.S. was supported by grants from the Deutsche Forschungsgemeinschaft (DFG; SCHL2276/2-1, 450149205-TRR333/1). We thank Frank Telge and Wolfgang Neumann for their help with constructing and producing the Slot Dot Blotting devices.

## References

1. Kerr, J.A., Patton, G.C., Cini, K.I., Abate, Y.H., Abbas, N., Abd Al Magied, A.H.A., Abd ElHafeez, S., Abd-Elsalam, S., Abdollahi, A., Abdoun, M., et al. (2025). Global, regional, and national prevalence of child and adolescent overweight and obesity, 1990-2021, with forecasts to 2050: a forecasting study for the Global Burden of Disease Study 2021. Lancet 405, 785–812. 10.1016/S0140-6736(25)00397-6.

2. Afshin, A., Forouzanfar, M.H., Reitsma, M.B., Sur, P., Estep, K., Lee, A., Marczak, L., Mokdad, A.H., Moradi-Lakeh, M., Naghavi, M., et al. (2017). Health Effects of Overweight and Obesity in 195 Countries over 25 Years. New Engl J Med 377, 13–27. 10.1056/NEJMoa1614362.

3. Heymsfield, S.B., and Wadden, T.A. (2017). Mechanisms, Pathophysiology, and Management of Obesity. New Engl J Med 376, 254–266. 10.1056/NEJMra1514009.

4. Halder, S.K., and Melkani, G.C. (2025). The Interplay of Genetic Predisposition, Circadian Misalignment, and Metabolic Regulation in Obesity. Curr Obes Rep 14. ARTN 21 10.1007/s13679-025-00613-3.

5. McCarthy, M.I. (2010). GENOMIC MEDICINE Genomics, Type 2 Diabetes, and Obesity. New Engl J Med 363, 2339–2350. DOI 10.1056/NEJMra0906948.

6. Farooqi, I.S. (2022). Monogenic Obesity Syndromes Provide Insights Into the Hypothalamic Regulation of Appetite and Associated Behaviors. Biol Psychiat 91, 856–859. 10.1016/j.biopsych.2022.01.018.

7. Kuenzel, R., Faust, H., Bundalian, L., Blueher, M., Jasaszwili, M., Kirstein, A., Kobelt, A., Koerner, A., Popp, D., Wenzel, E., et al. (2025). Detecting monogenic obesity: a systematic exome-wide workup of over 500 individuals. Int J Obesity. 10.1038/s41366-025-01819-0.

8. Barbosa, B.F., de Moraes, F.C.A., Barbosa, C.B., Santos, P.T.K.P., da Silva, I.P., da Silva, B.A.A., Barros, J.C.M., Burbano, R.M.R., dos Santos, N.P.C., and Fernandes, M.R. (2023). Efficacy and Safety of Setmelanotide, a Melanocortin-4 Receptor Agonist, for Obese Patients: A Systematic Review and Meta-Analysis. J Pers Med 13. ARTN 1460 10.3390/jpm13101460.

9. Argente, J., Verge, C.F., Okorie, U., Fennoy, I., Kelsey, M.M., Cokkinias, C., Scimia, C., Lee, H.M., and Farooqi, I.S. (2025). Setmelanotide in patients aged 2-5 years with rare MC4R pathway-associated obesity (VENTURE): a 1 year, open-label multicenter, phase 3 trial. Lancet Diabetes Endo 13, 29–37. 10.1016/S2213-8587(24)00273-0.

10. Morandi, A., Fornari, E., Corradi, M., Umano, G.R., Olivieri, F., Piona, C., Maguolo, A., Panzeri, C., Emiliani, F., Cirillo, G., et al. (2024). Variant reclassification over time decreases the level of diagnostic uncertainty in monogenic obesity: Experience from two centres. Pediatric Obesity 19. 10.1111/ijpo.13183.

11. Tamaroff, J., Williamson, D., Slaughter, J.C., Xu, M., Srivastava, G., and Shoemaker, A.H. (2023). Prevalence of genetic causes of obesity in clinical practice. Obes Sci Pract 9, 508–515. 10.1002/osp4.671.

12. Baldini, G., and Phelan, K.D. (2019). The melanocortin pathway and control of appetite-progress and therapeutic implications. J Endocrinol 241, R1–R33. 10.1530/JOE-18-0596.

13. Farooqi, I.S., Jebb, S.A., Langmack, G., Lawrence, E., Cheetham, C.H., Prentice, A.M., Hughes, I.A., McCamish, M.A., and O’Rahilly, S. (1999). Effects of recombinant leptin therapy in a child with congenital leptin deficiency. N Engl J Med 341, 879–884. 10.1056/NEJM199909163411204.

14. Farooqi, I.S., Wangensteen, T., Collins, S., Kimber, W., Matarese, G., Keogh, J.M., Lank, E., Bottomley, B., Lopez-Fernandez, J., Ferraz-Amaro, I., et al. (2007). Clinical and molecular genetic spectrum of congenital deficiency of the leptin receptor. N Engl J Med 356, 237–247. 10.1056/NEJMoa063988.

15. Clement, K., Vaisse, C., Lahlou, N., Cabrol, S., Pelloux, V., Cassuto, D., Gourmelen, M., Dina, C., Chambaz, J., Lacorte, J.M., et al. (1998). A mutation in the human leptin receptor gene causes obesity and pituitary dysfunction. Nature 392, 398–401. 10.1038/32911.

16. Montague, C.T., Farooqi, I.S., Whitehead, J.P., Soos, M.A., Rau, H., Wareham, N.J., Sewter, C.P., Digby, J.E., Mohammed, S.N., Hurst, J.A., et al. (1997). Congenital leptin deficiency is associated with severe early-onset obesity in humans. Nature 387, 903–908. 10.1038/43185.

17. Koerber-Rosso, I., Brandt, S., von Schnurbein, J., Fischer-Posovszky, P., Hoegel, J., Rabenstein, H., Siebert, R., and Wabitsch, M. (2021). A fresh look to the phenotype in mono-allelic likely pathogenic variants of the leptin and the leptin receptor gene. Mol Cell Pediatr 8, 10. 10.1186/s40348-021-00119-7.

18. Ernst, M.B., Wunderlich, C.M., Hess, S., Paehler, M., Mesaros, A., Koralov, S.B., Kleinridders, A., Husch, A., Münzberg, H., Hampel, B., et al. (2009). Enhanced Stat3 Activation in POMC Neurons Provokes Negative Feedback Inhibition of Leptin and Insulin Signaling in Obesity. J Neurosci 29, 11582–11593. 10.1523/Jneurosci.5712-08.2009.

19. Voigtmann, F., Wolf, P., Landgraf, K., Stein, R., Kratzsch, J., Schmitz, S., Abou Jamra, R., Bluher, M., Meiler, J., Beck-Sickinger, A.G., et al. (2021). Identification of a novel leptin receptor (LEPR) variant and proof of functional relevance directing treatment decisions in patients with morbid obesity. Metabolism 116, 154438. 10.1016/j.metabol.2020.154438.

20. von Schnurbein, J., Zorn, S., Nunziata, A., Brandt, S., Moepps, B., Funcke, J.B., Hussain, K., Farooqi, I.S., Fischer-Posovszky, P., and Wabitsch, M. (2024). Classification of Congenital Leptin Deficiency. J Clin Endocrinol Metab 109, 2602–2616. 10.1210/clinem/dgae149.

21. Saxton, R.A., Caveney, N.A., Moya-Garzon, M.D., Householder, K.D., Rodriguez, G.E., Burdsall, K.A., Long, J.Z., and Garcia, K.C. (2023). Structural insights into the mechanism of leptin receptor activation. Nat Commun 14, 1797. 10.1038/s41467-023-37169-6.

22. Park, H.K., and Ahima, R.S. (2014). Leptin signaling. F1000Prime Rep 6, 73. 10.12703/P6-73.

23. Gong, Y., Ishida-Takahashi, R., Villanueva, E.C., Fingar, D.C., Munzberg, H., and Myers, M.G., Jr. (2007). The long form of the leptin receptor regulates STAT5 and ribosomal protein S6 via alternate mechanisms. J Biol Chem 282, 31019–31027. 10.1074/jbc.M702838200.

24. Funcke, J.B., Moepps, B., Roos, J., von Schnurbein, J., Verstraete, K., Fröhlich-Reiterer, E., Kohlsdorf, K., Nunziata, A., Brandt, S., Tsirigotaki, A., et al. (2023). Rare Antagonistic Leptin Variants and Severe, Early-Onset Obesity. New Engl J Med 388, 2253–2261. 10.1056/NEJMoa2204041.

25. Mazen, I.H., El-Gammal, M.A., Elaidy, A.A., Anwar, G.M., Ashaat, E.A., Abdel-Ghafar, S.F., and Abdel-Hamid, M.S. (2023). Congenital leptin and leptin receptor deficiencies in nine new families: identification of six novel variants and review of literature. Mol Genet Genomics 298, 919–929. 10.1007/s00438-023-02025-1.

26. Kimber, W., Peelman, F., Prieur, X., Wangensteen, T., O’Rahilly, S., Tavernier, J., and Farooqi, I.S. (2008). Functional characterization of naturally occurring pathogenic mutations in the human leptin receptor. Endocrinology 149, 6043–6052. 10.1210/en.2008-0544.

27. Funcke, J.B., Moepps, B., Roos, J., von Schnurbein, J., Verstraete, K., Frohlich-Reiterer, E., Kohlsdorf, K., Nunziata, A., Brandt, S., Tsirigotaki, A., et al. (2023). Rare Antagonistic Leptin Variants and Severe, Early-Onset Obesity. N Engl J Med 388, 2253–2261. 10.1056/NEJMoa2204041.

28. Kimber, W., Peelman, F., Prieur, X., Wangensteen, T., O’Rahilly, S., Tavernier, J., and Farooqi, I.S. (2008). Functional Characterization of Naturally Occurring Pathogenic Mutations in the Human Leptin Receptor. Endocrinology 149, 6043–6052. 10.1210/en.2008-0544.

29. Delplanque, J., Le Collen, L., Loiselle, H., Leloire, A., Toussaint, B., Vaillant, E., Charpentier, G., Franc, S., Balkau, B., Marre, M., et al. (2024). Monoallelic pathogenic variants in LEPR do not cause obesity. Am J Hum Genet 111, 2668–2674. 10.1016/j.ajhg.2024.10.014.

30. Evans, M.C., Lord, R.A., and Anderson, G.M. (2021). Multiple Leptin Signalling Pathways in the Control of Metabolism and Fertility: A Means to Different Ends? Int J Mol Sci 22. 10.3390/ijms22179210.

31. Allison, M.B., and Myers, M.G., Jr. (2014). 20 years of leptin: connecting leptin signaling to biological function. J Endocrinol 223, T25–35. 10.1530/JOE-14-0404.

32. Liu, H., Du, T., Li, C., and Yang, G. (2021). STAT3 phosphorylation in central leptin resistance. Nutr Metab (Lond) 18, 39. 10.1186/s12986-021-00569-w.

33. Spielmann, M., and Kircher, M. (2022). Computational and experimental methods for classifying variants of unknown clinical significance. Csh Mol Case Stud 8. ARTN a006196 10.1101/mcs.a006196.

34. Töpf, A., Cox, D., Zaharieva, I.T., Di Leo, V., Sarparanta, J., Jonson, P.H., Sealy, I.M., Smolnikov, A., White, R.J., Vihola, A., et al. (2024). Digenic inheritance involving a muscle-specific protein kinase and the giant titin protein causes a skeletal muscle myopathy. Nat Genet 56. 10.1038/s41588-023-01651-0.

35. Bisello, G., and Bertoldi, M. (2022). Compound Heterozygosis in AADC Deficiency and Its Complex Phenotype in Terms of AADC Protein Population. International Journal of Molecular Sciences 23. ARTN 11238 10.3390/ijms231911238.

36. Ferrari, L., Mangano, E., Bonati, M.T., Monterosso, I., Capitanio, D., Chiappori, F., Brambilla, I., Gelfi, C., Battaglia, C., Bordoni, R., and Riva, P. (2020). Digenic inheritance of subclinical variants in Noonan Syndrome patients: an alternative pathogenic model? Eur J Hum Genet 28, 1432–1445. 10.1038/s41431-020-0658-0.

37. Sket, R., Kotnik, P., Bizjan, B.J., Kocen, V., Mlinaric, M., Tesovnik, T., Debeljak, M., Battelino, T., and Kovac, J. (2022). Heterozygous Genetic Variants in Autosomal Recessive Genes of the Leptin-Melanocortin Signalling Pathway Are Associated With the Development of Childhood Obesity. Front Endocrinol 13. ARTN 832911 10.3389/fendo.2022.832911.

38. Courbage, S., Poitou, C., Le Beyec-Le Bihan, J., Karsenty, A., Lemale, J., Pelloux, V., Lacorte, J.M., Carel, J.C., Lecomte, N., Storey, C., et al. (2021). Implication of Heterozygous Variants in Genes of the Leptin-Melanocortin Pathway in Severe Obesity. J Clin Endocr Metab 106, 2991–3006. 10.1210/clinem/dgab404.

39. González-Rodríguez, L., González, L.M., García-Herráiz, A., Mota-Zamorano, S., Flores, I., and Gervasini, G. (2025). Association of genetic variation in the leptin-melanocortin system with drive for thinness in patients with eating disorders: A pilot study. Gene 949. ARTN 149364 10.1016/j.gene.2025.149364.

40. Farooqi, I.S., Wangensteen, T., Collins, S., Kimber, W., Matarese, G., Keogh, J.M., Lank, E., Bottomley, B., Lopez-Fernandez, J., Ferraz-Amaro, I., et al. (2007). Clinical and molecular genetic spectrum of congenital deficiency of the leptin receptor. New Engl J Med 356, 237–247. DOI 10.1056/NEJMoa063988.

41. Gan, L.X., Guo, K.Y., Cremona, M.L., McGraw, T.E., Leibel, R.L., and Zhang, Y.Y. (2012). TNF-α Up-Regulates Protein Level and Cell Surface Expression of the Leptin Receptor by Stimulating Its Export via a PKC-Dependent Mechanism. Endocrinology 153, 5821–5833. 10.1210/en.2012-1510.

42. Bacart, J., Leloire, A., Levoye, A., Froguel, P., Jockers, R., and Couturier, C. (2010). Evidence for leptin receptor isoforms heteromerization at the cell surface. Febs Lett 584, 2213–2217. 10.1016/j.febslet.2010.03.033.

43. Barr, V.A., Lane, K., and Taylor, S.I. (1999). Subcellular localization and internalization of the four human leptin receptor isoforms. J Biol Chem 274, 21416–21424. 10.1074/jbc.274.30.21416.

44. Gorska, E., Popko, K., Stelmaszczyk-Emmel, A., Ciepiela, O., Kucharska, A., and Wasik, M. (2010). Leptin receptors. Eur J Med Res 15 *Suppl 2*, 50–54. 10.1186/2047-783x-15-s2-50.

45. Bjorbaek, C., Uotani, S., da Silva, B., and Flier, J.S. (1997). Divergent signaling capacities of the long and short isoforms of the leptin receptor. Journal of Biological Chemistry 272, 32686–32695. DOI 10.1074/jbc.272.51.32686.

46. Banks, W.A. (2001). Leptin transport across the blood-brain barrier: implications for the cause and treatment of obesity. Curr Pharm Des 7, 125–133. 10.2174/1381612013398310.

47. Landt, M., Parvin, C.A., and Wong, M. (2000). Leptin in cerebrospinal fluid from children: correlation with plasma leptin, sexual dimorphism, and lack of protein binding. Clin Chem 46, 854–858.

