## Supplemental Figures for "A simple, cost efficient assay for assessing the functional impact of single and multi-gene variant combinations"

### Fig.S1

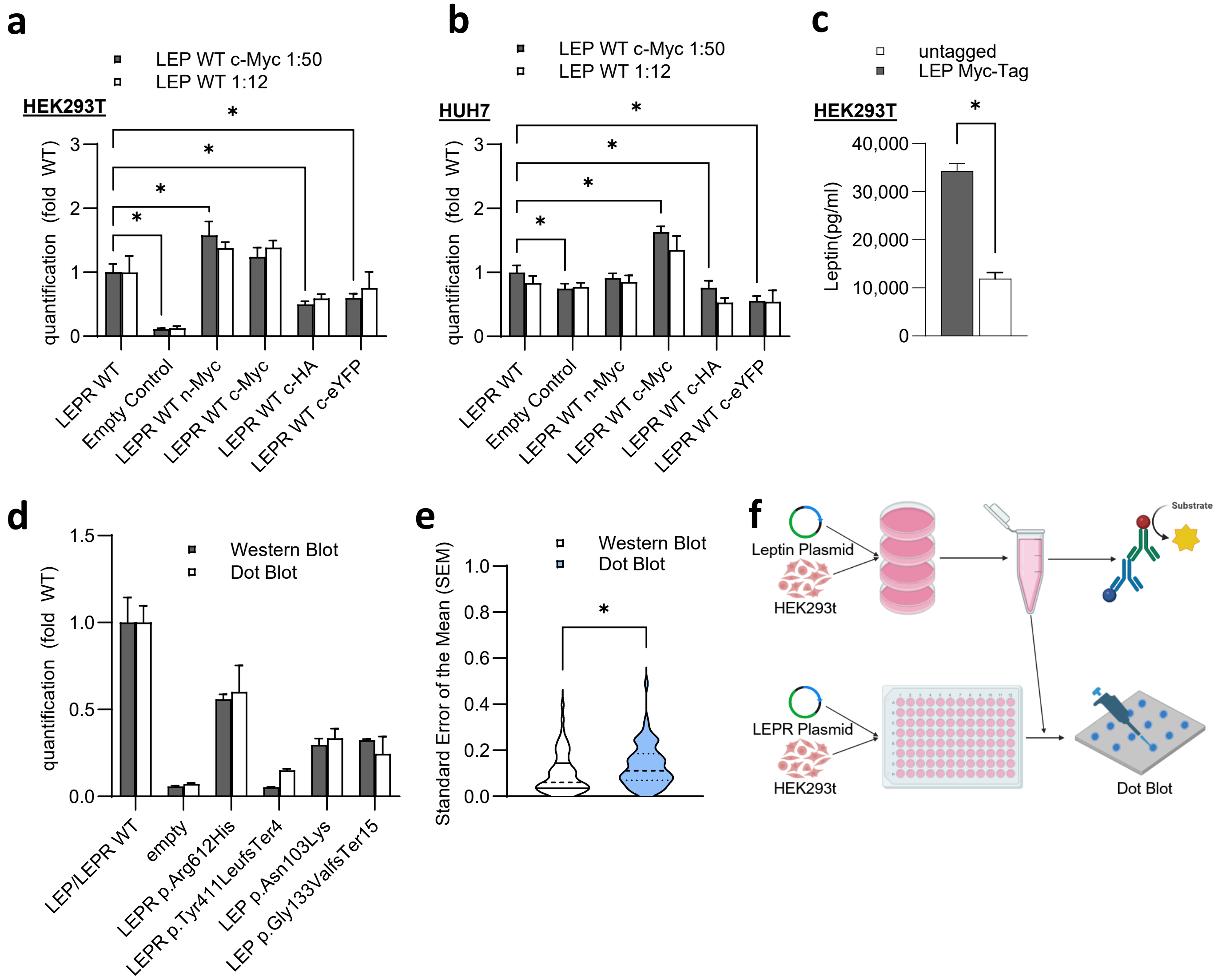


**Fig.S1: Assay design and validation for functional characterization of *LEP* and *LEPR* variants.** (A) Comparison of phosphoStat3 DB data for *LEPR* WT untagged, Empty Control, *LEPR* WT n-Myc, *LEPR* WT c-Myc, *LEPR* WT c-HA and *LEPR* WT c-eYFP with n=4 for both *LEP* WT untagged and *LEP* WT c-Myc for every variant in HEK293t cells. (B) Comparison of phosphoStat3 DB data for LEPR WT untagged, Empty Control, *LEPR* WT n-Myc, *LEPR* WT c-Myc, *LEPR* WT c-HA and *LEPR* WT c-eYFP with n=4 for both *LEP* WT untagged and *LEP* WT c-Myc for every variant in HuH-7 cells. (C) Amount of leptin Wild Type c-myc and untagged (in pg/ml) in Supernatants of HEK293T cells measured with a leptin-ELISA. (D) Comparison of a Western Blot and a Dot Blot for *LEP* and *LEPR* variant lysates generated in HEK293T cells and the respective level of phosphorylated STAT3 compared to wild type levels. (E) Comparison of the SEM of Dot Blot and Western Blot for the same variants. (F) The process of generating leptin supernatants protein lysates of variant combinations for Dot Blot. The picture was generated using BioRender.

### Fig.S2

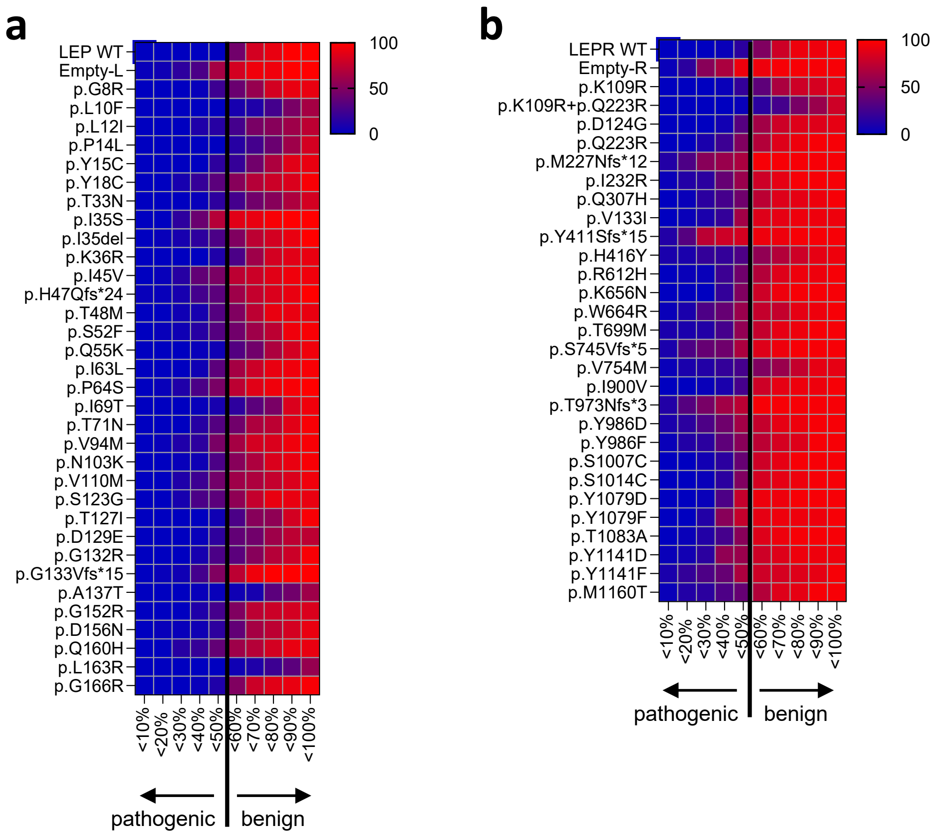


**Fig.S2: Screening of common Leptin and Leptin Receptor variants and their compound-heterozygous-like combination.** (A) A heat map presentation of the percentage of the compound-heterozygous-like combinations of *LEP* variants below different thresholds between <10% and <100% of wild type signalling levels. (B) A heat map presentation of the percentage of the compound-heterozygous-like combinations of *LEPR* variants below different thresholds between <10% and <100% of wild type signalling levels.
